# Physiological Profiles of Male and Female CrossFit^®^ Athletes

**DOI:** 10.1101/2023.10.11.561828

**Authors:** Deni Hodžić, Gommaar D’Hulst, Rahel Leuenberger, Janik Arnet, Elena Westerhuis, Ralf Roth, Arno Schmidt-Trucksäss, Raphael Knaier, Jonathan Wagner

## Abstract

**Objective:** To 1) establish extensive physiological profiles of highly-trained CrossFit athletes using gold-standard tests and 2) investigate which physiological markers best correlate with CrossFit^®^ Open performance.

**Methods:** This study encompassed sixty participants (30 males and 30 females), all within the top 5% of the CrossFit^®^ Open, including 7 CrossFit^®^ Semi-finalists and 3 Games finalists. Isokinetic dynamometers were employed to measure maximum isometric and isokinetic leg and trunk strength. Countermovement jump height and maximum isometric mid-thigh pull strength were assessed on a force plate. V□O_2peak_ was measured by a cardiopulmonary exercise test, while critical power and W’ were evaluated during a 3-minute all-out test, both on a cycle ergometer.

**Results:** Male and female athletes’ median (IQR) V□O_2peak_ was 4.64 (4.43, 4.80) and 3.21 (3.10, 3.29) (L·min^-1^), critical power 314.5 (285.9, 343.6) and 221.3 (200.9, 238.9) (W) and mid-thigh pull 3158 (2690, 3462) and 2035 (1728, 2347) (N). Linear regression analysis shows strong evidence for associations between different anthropometric variables and CrossFit^®^ Open performance in men and women, whereas for markers of cardiorespiratory fitness such as V□O_2peak_ this was only true for women but not men. Conventional laboratory evaluations of strength, however, manifest minimal evidence for associations with Open performance across both sexes.

**Conclusions:** This study provides the first detailed insights into the physiology of high-performing CrossFit^®^ athletes and informs training optimization. Further the results emphasize the advantage of athletes with shorter limbs and suggests potential modifications to Open workout designs to level the playing field for athletes across different anthropometrics characteristics.

## INTRODUCTION

CrossFit^®^, a sport that pushes the boundaries of human physiological capacity and strength through concurrent training, presents a unique arena to test physical fitness on a global scale. Since 2011, CrossFit^®^ Inc. has hosted a worldwide Open competition that invites hundreds of thousands of athletes to perform the same workouts over 3 to 5 weekends, culminating in the most extensive fitness test globally.^1^

In an effort to predict performance in this open competition, numerous studies have adopted basic physiological profiling and endurance testing, predominantly using regression analysis.^2–5^ However, these investigations have not targeted the upper echelon athletes participating in the CrossFit^®^ Games, leaving a significant gap in our understanding of what defines top-tier CrossFit^®^ performance. Furthermore, while extensive physiological profiling has been conducted for sports like cycling^6,7^ and weightlifting^8–10^, such comprehensive profiling is largely absent in CrossFit^®^. This lack of profiling constitutes a significant gap in knowledge, especially considering the increasing popularity and competitive nature of the sport.

The primary aim of this study is to establish a comprehensive physiological, anthropometric, and strength profile of highly trained CrossFit^®^ athletes—those who rank in the top 5% of the CrossFit^®^ Open, including Semi-finalists and CrossFit^®^ Games competitors. Additionally, we aim to identify which conventional laboratory-based tests most accurately predict performance in the CrossFit^®^ Open among this elite cohort.

In providing a detailed profile of top-performing CrossFit^®^ athletes and identifying key performance predictors, this study provides valuable insights to the growing body of knowledge surrounding the sport. More importantly, these findings will serve as a valuable tool for coaches and athletes alike, offering a strategic roadmap to excel in this unique and demanding discipline.

## METHODS

### Study design

The PhyPCross study was conducted between April 2021 and July 2022 in the laboratories of the Department of Sport, Exercise and Health at the University of Basel, Switzerland. All participants gave written informed consent before inclusion, and the study was reviewed and approved by the cantonal ethics committee of Northwest and Central Switzerland (EKNZ-2021-00650). This cross-sectional single-center study involved one examination date, lasting approximately three hours in total. The exact measurement procedure is shown in Figure 1.

**Figure 1.**
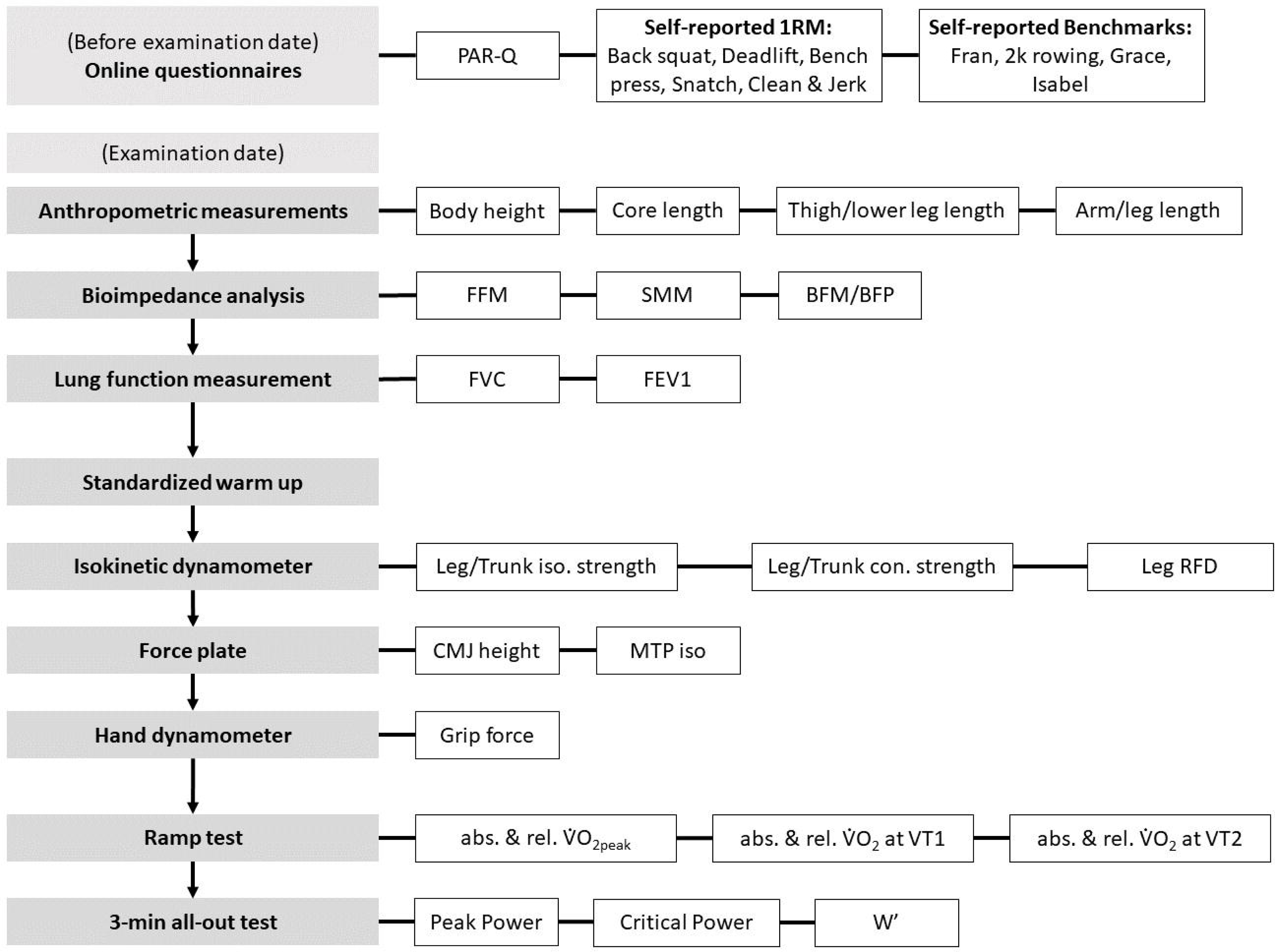
data acquisition scheme. Abbreviations: PAR-Q, Physical Activity Readiness Questionnaire; BIA, bioimpedance analysis; FFM, fat free mass; SMM, skeletal muscle mass; BFM, body fat mass; BFP, body fat percentage; FVC, forced vital capacity; FEV1, forced expiratory volume after one second; iso, isometric; con, concentric; RFD, rate of force development; CMJ, countermovement jump; MTP, mid-thigh pull; abs, absolute; rel, relative; VDO_2peak_, peak oxygen consumption; VT1, first ventilatory threshold; VT2, second ventilatory threshold; W’, work capacity above critical power.

### Participants

Inclusion criteria were age between 18 and 40 years, a top 5% finish at CrossFit^®^ Opens between 2019-2022 and currently training for the next competitive season. Exclusion criteria included febrile infection within the last 14 days, known cardiovascular disease, diabetes mellitus type 1 or type 2, hypertension (systolic/diastolic blood pressure >160 mmHg/ >100 mmHg), or participation in another clinical trial in the past four weeks.

Participants were recruited through outreach in and outside the University of Basel, direct inquiries from CrossFit^®^ boxes, and collaboration with the ETH Zurich and the scientific social media site “WODSience”. The detailed recruitment procedure is shown in Figure 2.

**Figure 2.**
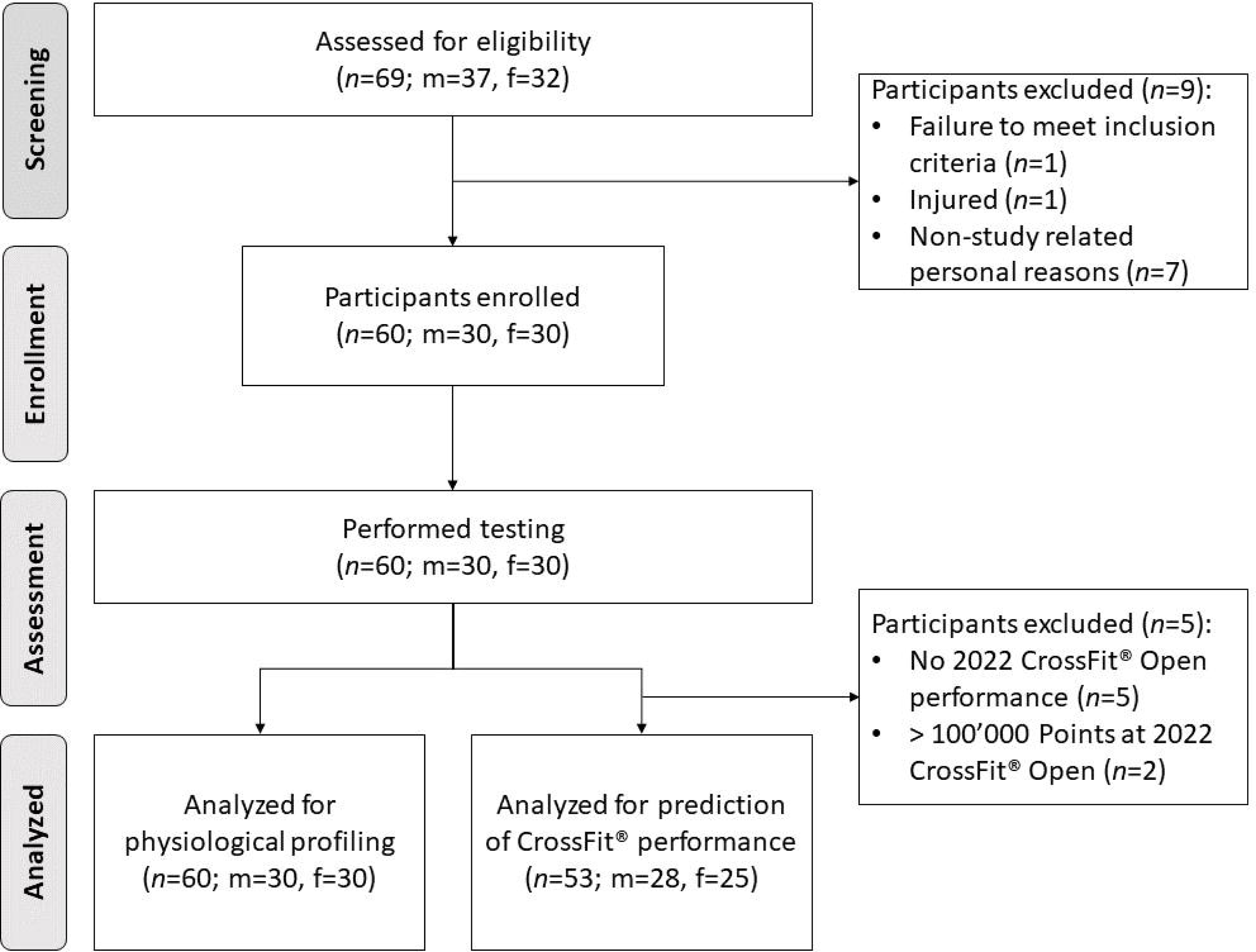
Flow diagram showing inclusion and exclusion of participants. Abbreviations: n, number of observations; m, male; f, female.

All participants were considered highly-trained and divided into either well-trained or elite athletes based on their performance at the 2022 CrossFit^®^ Open. Elite athletes were defined as participants qualifying for the semi-finals, whereas well-trained athletes were defined as all the other participants.

### Acquisition of participant characteristics

Two online questionnaires were completed before the examination date. All participants filled out the Physical Activity Readiness Questionnaire (PAR-Q) ^11^ and if abnormalities were detected, a preliminary medical examination was performed. Additionally, the self-reported one repetition maximum (1RM) values of “Back Squat”, “Deadlift”, “Bench Press”, “Snatch”, and “Clean and Jerk” and the benchmarks “Fran”, “2’000m Rowing”, “Grace”, and “Isabel” were asked. After a detailed explanation of the testing procedure, anthropometric data were measured. Body mass and composition were measured using a four-segment bioelectrical impedance analysis (Inbody 720; Inbody Co. Ltd., Seoul, South Korea). Participants were instructed not to consume caffeinated beverages for four hours and food for two hours prior to testing. More detailed information on the acquisition of participant characteristics is provided in the supplement.

### Strength testing

All strength assessments were performed using the gold-standard method isokinetic dynamometers (IsoMed 2000, D&R Ferstl, Hemau, Germany). Before strength testing, all participants performed a standardized warm-up program. For all tests, participants were verbally supported to ensure maximal effort.

### Leg strength test

The participants were fixed in a standardized upright sitting position at both shoulders and with a belt around their hip. For the measurement of isometric leg strength, the knee angle was set at 140° and the hip angle at 135°. The hands were placed on handles at waist level. The feet were positioned flat on the support surface, with the heels on level with the seat. For the measurement of isokinetic leg strength, the knee angle was adjusted from 90° to 110°. The foot support surface moved at a constant speed of 120 mm·s^-1^ between the two endpoints.

### Trunk strength test

Participants were seated on the device with a hip angle of 85° and a knee angle of 90°. In this position, the lower legs, thighs, and shoulder girdle were fixed. Hands were placed on two grips in front of the chest near the collarbone.

Isokinetic trunk strength measurement during concentric flexion and extension was performed at a movement speed of 60°·s^-1^. The range of motion ranged from a hip angle of 55° to 115°. The rotation point of the device was verified with a laser oriented to the upper edge of the iliac crest.

### Countermovement Jump

A countermovement jump on the force plate (Leonardo Mechanography, Novotec Medical GmbH, Pforzheim, Germany) was performed to determine lower extremities explosivity. The countermovement depth could be chosen individually and the hands were placed on the hips. Participants were instructed to jump as high as possible. When landing, the legs ought to remain extended. Jump height was calculated based on the measured flight time. As previous studies show, jump mechanography is reliable for assessing lower extremities’ musculoskeletal function.^12,13^

### Mid-thigh pull strength testing

The maximum isometric force of the lower extremities was determined in the mid-thigh pull position on the force plate. The starting position was with the fixed bar at mid-thigh and a knee and hip angle of 140-160°. The grip position was outside the thighs. The starting position was maintained throughout the force measurement. Participants were instructed to press the legs into the ground as “quickly” and as “strongly” as possible for at least six seconds.

### Grip force testing

The maximum grip force of the dominant hand was measured using a hand-held dynamometer (Leonardo Mechanograph GF, Novotec Medical GmbH, Pforzheim, Germany). The starting position was upright, with the tested arm allowed to hang loosely. The forearm was in a neutral position, the wrist in a 15° extension. From this position, the task was to apply pressure to the device as “strongly” as possible while maintaining the body posture. Span width was individualized.^14^

### Cardiopulmonary exercise testing

A cardiopulmonary exercise test using a ramp protocol was performed on a cycle ergometer (Excalibur Sport, Lode BV, Groningen, The Netherlands) with simultaneous breath-by-breath gas analysis (MetaMax 3B and MetaSoft^®^ Studio software, version 5.8.5 SR+, Cortex, Leipzig, Germany). V□O_2peak_ was defined as the highest 30-second average of V□O_2_ at any point during the test.

### Ramp test

After a 5-min warm-up at 100 W for males and 50 W for females, the workload increased linearly with 30 W per min until the incremental test was terminated. The test was aborted either by the participant’s exhaustion or by the researcher if, after a single warning, the cadence could no longer be kept above 60 rpm. During the test, a 12-lead ECG (Custo Med GmbH, Ottobrunn, Germany) was derived. The heart rate was measured using a Polar chest strap (Polar Electro Oy, Kempele, Finland) during the entire test period and recorded directly via the MetaSoft^®^ Studio program

### 3-min all-out test

Immediately before the 3-min all-out test participants performed a warm-up phase of three minutes without resistance. During the last ten seconds of the warm-up phase, participants were instructed to increase their cadence to approximately 110-120 rpm. The maximum possible power was to be achieved at the beginning of the 3-minute test phase. Subsequently, the highest possible cadence had to be maintained against a linear resistance for the entire duration of the test. For this purpose, the alpha value (α) was calculated as suggested by Vanhatalo et al.^16^ No information about the remaining time and cadence was given to prevent pacing. The critical power is calculated from the average power during the last 30 seconds of the test. To estimate the W’, the power-time integral was calculated over the critical power.^16,17^

### Statistical Analysis

Participants have a publicly accessible online profile on the official website of the CrossFit^®^ Open, listing the ranking history of the respective competition phase from previous years. Performance data from the 2022 CrossFit^®^ Open were recorded for all participants. In this study, the “Points” at the 2022 CrossFit^®^ Open were considered, which is the sum of the ranks in the individual workouts and not the “Rank”, which is the “Points” in comparison with all the other competing athletes.

Residual diagnostics were used to evaluate the satisfaction of model assumptions for t-test and linear regression models. Robust standard errors were used where heteroscedasticity was detected. Physiological profiles are shown as percentiles of all athletes and both medians for well-trained and elite athletes. An independent t-test was used to compare mean differences between well-trained and elite athletes. Standard deviations, 95% confidence intervals and p-values were calculated. To investigate the association between single physiological parameters and CrossFit^®^ Open performance, linear regression analyses with number of points at the 2022 Open as the dependent variable and the single physiological parameters as independent variables were calculated. R statistical Software (version 4.3.0) was used for all analyses.

## RESULTS

### Physiological Profiling

In this study, physiological profiling was conducted on 60 participants, 30 males and 30 females. The details of the participants’ general characteristics and their performances in the 2022 CrossFit^®^ Open are presented in Table 1. The athletes were also asked to provide their self-reported results for their CrossFit^®^ benchmark performances as part of a questionnaire, of which the data is also presented in Table 1.

**Table 1.** Descriptive Characteristics of study participants.

|  | All | Men | Women |
| --- | --- | --- | --- |
| **General characteristics** |  |  |  |
| n | 60 | 30 | 30 |
| Age (years) | 28.3 (4.4) | 27.6 (4.1) | 29.1 (4.7) |
| Height (cm) | 170.7 (8.3) | 176.3 (6.3) | 165.5 (6.5) |
| Body mass (kg) | 75.6 (12.3) | 85.9 (7.7) | 65.9 (6.5) |
| Body mass index (kg·m<sup>-2</sup>) | 25.8 (2.5) | 27.6 (1.5) | 24.1 (2.0) |
| **Benchmarks** |  |  |  |
| Back Squat (kg) | 149.6 (35.7) (n=59) | 178.3 (17.7) (n=29) | 115.0 (13.8) (n=30) |
| Deadlift (kg) | 179.1 (41.1) (n=59) | 211.6 (19.1) (n=29) | 138.3 (16.1) (n=30) |
| Bench Press (kg) | 105.0 (31.4) (n=58) | 127.9 (17.9) (n=30) | 72.5 (9.2) (n=28) |
| Snatch (kg) | 91.4 (20.6) (n=57) | 106.8 (10.6) (n=28) | 71.3 (9.7) (n=27) |
| Clean and Jerk (kg) | 113.8 (24.3) (n=57) | 132.2 (11.9) (n=28) | 88.6 (10.1) (n=27) |
| Fran (s) | 180.7 (46.6) (n=32) | 157.3 (31.4) (n=20) | 217.1 (43.6) (n=12) |
| 2k row (s) | 426.7 (64.0) (n=34) | 396.4 (58.3) (n=17) | 457.1 (54.3) (n=17) |
| Grace (s) | 128.0 (34.8) (n=28) | 118.8 (33.3) (n=19) | 147.4 (29.4) (n=9) |
| Isabel (s) | 141.0 (49.5) (n=23) | 119.6 (35.1) (n=14) | 174.3 (50.2) (n=9) |
| **2022 CrossFit® performance** |  |  |  |
| Open Participations | 55 | 29 | 26 |
| Opens Ranks | 5451 (11264) | 5664 (9415) | 5205 (18191) |
| Opens Points | 19505 (35364) | 20644 (31249) | 13071 (39544) |
| Quarters Participations | 36 | 18 | 18 |
| Quarters Ranks | 425 (368) | 421 (8870) | 429 (8557) |
| Quarters Points | 8713 (6226) | 344 (5548) | 392 (6833) |
| Semis Participations | 7 | 4 | 3 |
| Semis Ranks | 12 (8) | 14 (295) | 11 (346) |
| Semis Points | 316 (107) | 9 (123) | 5 (69) |
| Games Participations | 3 | 2 | 1 |
| Games Ranks | 26 (9) | 22 (584) | 35 (296) |
| Games Points | 488 (178) | 7 (141) | 0 (0) |
| Data are mean (standard deviation). |  |  |  |

Table 2 provides a comprehensive overview of the athletes’ physiological profiles including different percentiles for a more detailed analysis. When comparing well-trained male to elite male athletes there was evidence for differences in relative V□O_2_ at the second ventilatory threshold (VT2). Similarly, among the female participants, there was evidence for differences between the two groups in the areas of body fat mass, isometric trunk flexion and concentric trunk flexion.

**Table 2.** Physiological profiles of highly trained athletes (left side). Group comparison of well-trained athletes versus elite athletes (right side).

| Physiologic parameter | Percentiles |  |  |  |  | Well-trained athletes | Elite athletes | Well-trained vs Elite athletes |  |
| --- | --- | --- | --- | --- | --- | --- | --- | --- | --- |
|  | 5<sup>th</sup> | 25<sup>th</sup> | 50<sup>th</sup> | 75<sup>th</sup> | 95<sup>th</sup> | Median | Median | Mean Difference (SD) | 95% CI |
| **MEN** |  |  |  |  |  |  |  |  |  |
| **Anthropometrics** |  |  |  |  |  |  |  |  |  |
| Height (cm) | 167.4 | 172.0 | 175.5 | 180.8 | 187.0 | 176.0 | 173.5 | 3.2 (2.3) | -2.2; 8.6 |
| Core length (cm) | 43.0 | 47.1 | 48.5 | 51.2 | 52.7 | 48.5 | 47.8 | 2.9 (2.2) | -3.5; 9.3 |
| Arm length (cm) | 52.4 | 57.0 | 58.0 | 60.0 | 63.6 | 58.3 | 57.0 | 2.8 (2.1) | -3.4; 8.9 |
| Leg length (cm) | 88.5 | 92.1 | 95.0 | 97.9 | 102.7 | 95.0 | 92.5 | 3.0 (2.5) | -4.0; 10.0 |
| Thigh length (cm) | 52.5 | 55.1 | 58.3 | 60.0 | 62.5 | 58.3 | 55.5 | 2.4 (2.5) | -5.1; 9.8 |
| Lower leg length (cm) | 33.5 | 35.5 | 37.0 | 39.6 | 42.1 | 37.0 | 37.0 | 0.6 (0.8) | -1.2; 2.4 |
| **Body composition** |  |  |  |  |  |  |  |  |  |
| Body mass (kg) | 74.1 | 81.5 | 85.5 | 90.8 | 100.3 | 86.5 | 82.1 | 4.5 (2.4) | -0.9; 9.8 |
| FFM (kg) | 64.7 | 73.0 | 77.6 | 80.1 | 90.6 | 77.6 | 75.4 | 2.4 (2.5) | -3.4; 8.2 |
| SMM (kg) | 37.2 | 42.2 | 44.9 | 46.5 | 52.7 | 44.9 | 43.8 | 1.3 (1.5) | -2.2; 4.7 |
| BFM (kg) | 5.4 | 6.5 | 8.2 | 10.7 | 14.2 | 8.6 | 6.7 | 2.1 (0.9) | 0.0; 4.2 |
| BFP (%) | 5.9 | 7.9 | 10.3 | 11.9 | 16.5 | 10.7 | 7.8 | 2.0 (1.2) | -0.7; 4.8 |
| Body mass index (kg·m<sup>-2</sup>) | 25.5 | 26.6 | 27.5 | 28.7 | 29.8 | 27.5 | 27.3 | 0.4 (0.7) | -1.5; 2.2 |
| **Endurance** |  |  |  |  |  |  |  |  |  |
| V̇O<sub>2peak</sub> (L·min<sup>-1</sup>) | 4.03 | 4.43 | 4.64 | 4.80 | 5.08 | 4.60 | 4.70 | -0.0 (0.2) | -0.4; 0.3 |
| V̇O<sub>2peak</sub> (mL·kg<sup>-1</sup>·min<sup>-1</sup>) | 48.0 | 51.2 | 53.7 | 56.2 | 59.1 | 53.3 | 56.3 | -3.4 (1.5) | -7.2; 0.4 |
| V̇O<sub>2</sub> at VT1 (mL·kg<sup>-1</sup>·min<sup>-1</sup>) | 27.3 | 31.1 | 33.4 | 36.6 | 44.1 | 32.5 | 37.9 | -6.2 (2.6) | -13.6; 1.1 |
| V̇O<sub>2</sub> at VT2 (mL·kg<sup>-1</sup>·min<sup>-1</sup>) | 42.8 | 46.5 | 48.2 | 49.8 | 53.1 | 47.7 | 51.5 | -3.9 (1.2) | -7.1; -0.7 |
| Peak Power (W) | 770.3 | 973.8 | 1067.5 | 1342.5 | 1704.9 | 1065.5 | 1163.0 | -2.1 (190.0) | -549.7; 545.4 |
| Critical Power (W) (n=29) | 245.2 | 285.9 | 314.5 | 343.6 | 381.2 | 314.5 | 334.4 | -32.1 (41.0) | -156.7; 92.5 |
| W' (kJ) | 9.3 | 15.4 | 20.9 | 23.1 | 26.0 | 20.9 | 19.6 | 3.4 (6.8) | -17.7; 24.5 |
| **Strength** |  |  |  |  |  |  |  |  |  |
| Leg iso. (N) | 3186 | 4116 | 4962 | 5416 | 6493 | 4842 | 5288 | -216.6 (585.1) | -1827.4; 1394.3 |
| Leg max con. [120 mm·s<sup>-1</sup>] (N) | 2479 | 3023 | 3516 | 3796 | 4541 | 3504 | 3570 | 96.8 (300.6) | -639.1; 832.7 |
| Trunk iso. ext. (N) | 3020 | 3625 | 4233 | 4840 | 5293 | 4489 | 3296 | 1173.7 (407.7) | 36.5; 2310.9 |
| Trunk iso. flex. (N) | 1368 | 1707 | 1961 | 2160 | 2551 | 1908 | 2211 | -136.8 (175.6) | -568.4; 294.8 |
| Trunk con. ext. (N) | 3339 | 3867 | 4193 | 4630 | 4976 | 4077 | 4297 | -19.0 (161.0) | -360.0; 322.0 |
| Trunk con. flex. (N) | 1378 | 1662 | 1772 | 2003 | 2322 | 1772 | 1792 | 25.2 (121.2) | -265.1; 315.6 |
| CMJ height (m) | 0.33 | 0.36 | 0.39 | 0.44 | 0.52 | 0.38 | 0.43 | -0.0 (0.0) | -0.1; 0.0 |
| max. MTP (N) | 1979 | 2690 | 3158 | 3462 | 3693 | 3184 | 2894 | 99.9 (351.5) | -913.9; 1113.7 |
| max GF (N) | 460 | 518 | 564 | 650 | 792 | 579 | 526 | 79.3 (24.3) | 28.7; 129.8 |
| **WOMEN** |  |  |  |  |  |  |  |  |  |
| **Anthropometrics** |  |  |  |  |  |  |  |  |  |
| Height (cm) | 155.0 | 163.0 | 166.0 | 169.5 | 175.0 | 166.0 | 167.0 | 1.3 (5.0) | -17.8; 20.4 |
| Core length (cm) | 40.7 | 43.0 | 44.5 | 48.0 | 49.6 | 45.0 | 42.0 | 1.7 (2.5) | -8.2; 11.5 |
| Arm length (cm) | 49.5 | 52.0 | 54.0 | 55.0 | 56.0 | 54.0 | 52.0 | 1.5 (1.2) | -2.9; 5.9 |
| Leg length (cm) | 84.2 | 87.0 | 90.8 | 93.0 | 96.6 | 91.0 | 90.0 | 1.9 (2.2) | -5.5; 9.3 |
| Thigh length (cm) | 50.5 | 53.0 | 55.0 | 57.4 | 61.0 | 55.0 | 52.0 | 2.2 (2.0) | -4.6; 9.1 |
| Lower leg length (cm) | 32.0 | 33.0 | 35.0 | 36.0 | 38.6 | 35.0 | 34.0 | -0.3 (1.5) | -5.9; 5.2 |
| **Body composition** |  |  |  |  |  |  |  |  |  |
| Body mass (kg) | 58.0 | 61.8 | 64.7 | 69.1 | 75.9 | 66.5 | 64.5 | 2.5 (1.6) | -0.9; 5.9 |
| FFM (kg) | 47.6 | 52.0 | 56.4 | 58.5 | 64.0 | 56.4 | 56.4 | 0.1 (1.8) | -4.4; 4.6 |
| SMM (kg) | 26.5 | 29.2 | 32.1 | 33.5 | 36.4 | 32.3 | 31.9 | -0.1 (1.1) | -2.8; 2.6 |
| BFM (kg) | 5.7 | 6.9 | 9.5 | 11.7 | 16.6 | 10.0 | 8.1 | 2.4 (0.9) | 0.3; 4.4 |
| BFP (%) | 9.6 | 10.8 | 13.7 | 18.7 | 23.4 | 13.8 | 12.5 | 3.0 (1.4) | -0.4; 6.3 |
| Body mass index (kg·m<sup>-2</sup>) | 21.2 | 22.6 | 24.2 | 25.5 | 27.6 | 24.3 | 23.1 | 0.5 (1.2) | -3.7; 4.6 |

### Predicting CrossFit^®^ Open Performance

Table 3 presents a linear regression analysis that correlates anthropometric, endurance, and strength parameters with CrossFit^®^ Open ranking in this highly trained CrossFit^®^ cohort. Notably, we found strong evidence for positive associations between various anthropometric parameters—including height, thigh length, and body weight— and Open performance. This correlation was especially pronounced among male participants, implying a stronger link between anthropometric factors and performance in men than in women. Table 3 further reveals interesting patterns regarding endurance and strength parameters. In women, variables such as relative V□O_2peak_, along with relative V□O_2_ at VT2, were predominantly negatively associated with Open rank, indicating a positive influence on performance. However, these variables had no significant impact on men’s Open performance. Critical power displayed a tendency towards a negative correlation with rank in both sexes, suggesting its beneficial influence on performance.

**Table 3.**
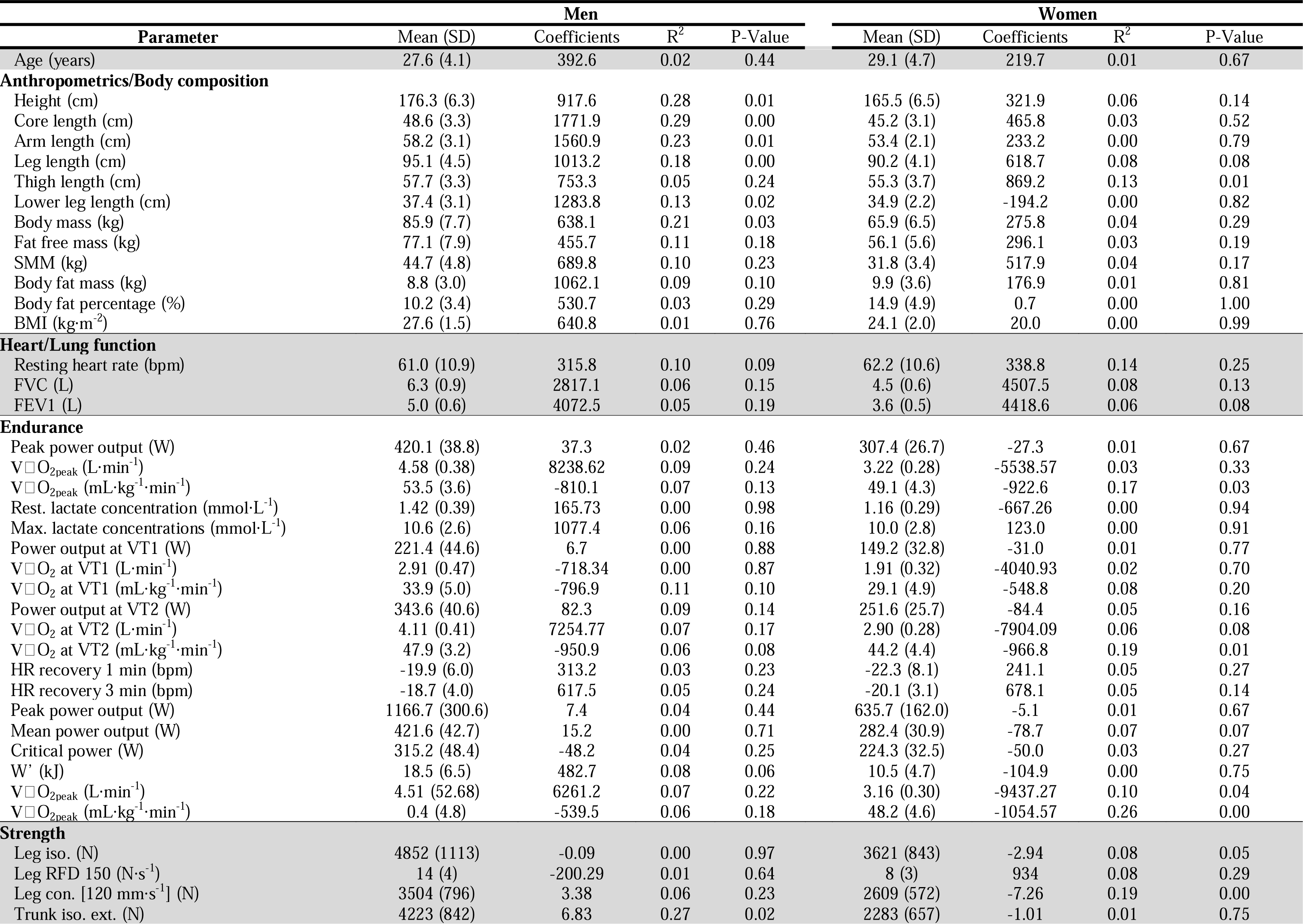
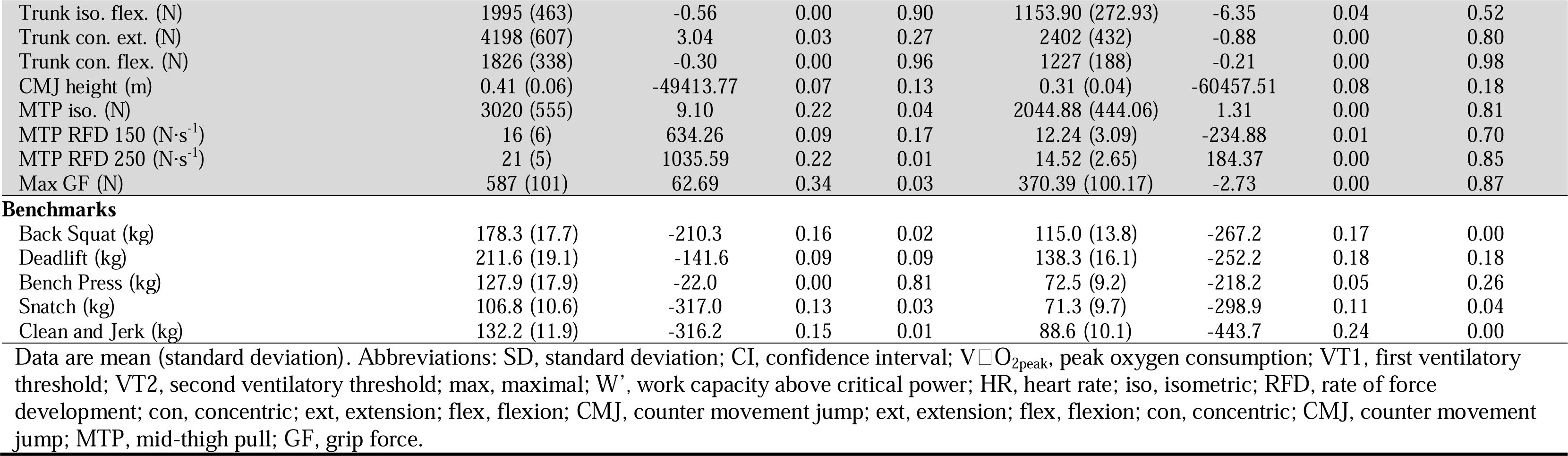
Associations between single parameters and CrossFit^®^ Open performance.

| Parameter | Men |  |  |  | Women |  |  |  |
| --- | --- | --- | --- | --- | --- | --- | --- | --- |
|  | Mean (SD) | Coefficients | R<sup>2</sup> | P-Value | Mean (SD) | Coefficients | R<sup>2</sup> | P-Value |
| Age (years) | 27.6 (4.1) | 392.6 | 0.02 | 0.44 | 29.1 (4.7) | 219.7 | 0.01 | 0.67 |
| **Anthropometrics/Body composition** |  |  |  |  |  |  |  |  |
| Height (cm) | 176.3 (6.3) | 917.6 | 0.28 | 0.01 | 165.5 (6.5) | 321.9 | 0.06 | 0.14 |
| Core length (cm) | 48.6 (3.3) | 1771.9 | 0.29 | 0.00 | 45.2 (3.1) | 465.8 | 0.03 | 0.52 |
| Arm length (cm) | 58.2 (3.1) | 1560.9 | 0.23 | 0.01 | 53.4 (2.1) | 233.2 | 0.00 | 0.79 |
| Leg length (cm) | 95.1 (4.5) | 1013.2 | 0.18 | 0.00 | 90.2 (4.1) | 618.7 | 0.08 | 0.08 |
| Thigh length (cm) | 57.7 (3.3) | 753.3 | 0.05 | 0.24 | 55.3 (3.7) | 869.2 | 0.13 | 0.01 |
| Lower leg length (cm) | 37.4 (3.1) | 1283.8 | 0.13 | 0.02 | 34.9 (2.2) | -194.2 | 0.00 | 0.82 |
| Body mass (kg) | 85.9 (7.7) | 638.1 | 0.21 | 0.03 | 65.9 (6.5) | 275.8 | 0.04 | 0.29 |
| Fat free mass (kg) | 77.1 (7.9) | 455.7 | 0.11 | 0.18 | 56.1 (5.6) | 296.1 | 0.03 | 0.19 |
| SMM (kg) | 44.7 (4.8) | 689.8 | 0.10 | 0.23 | 31.8 (3.4) | 517.9 | 0.04 | 0.17 |
| Body fat mass (kg) | 8.8 (3.0) | 1062.1 | 0.09 | 0.10 | 9.9 (3.6) | 176.9 | 0.01 | 0.81 |
| Body fat percentage (%) | 10.2 (3.4) | 530.7 | 0.03 | 0.29 | 14.9 (4.9) | 0.7 | 0.00 | 1.00 |
| BMI (kg·m<sup>-2</sup>) | 27.6 (1.5) | 640.8 | 0.01 | 0.76 | 24.1 (2.0) | 20.0 | 0.00 | 0.99 |
| **Heart/Lung function** |  |  |  |  |  |  |  |  |
| Resting heart rate (bpm) | 61.0 (10.9) | 315.8 | 0.10 | 0.09 | 62.2 (10.6) | 338.8 | 0.14 | 0.25 |
| FVC (L) | 6.3 (0.9) | 2817.1 | 0.06 | 0.15 | 4.5 (0.6) | 4507.5 | 0.08 | 0.13 |
| FEV1 (L) | 5.0 (0.6) | 4072.5 | 0.05 | 0.19 | 3.6 (0.5) | 4418.6 | 0.06 | 0.08 |
| **Endurance** |  |  |  |  |  |  |  |  |
| Peak power output (W) | 420.1 (38.8) | 37.3 | 0.02 | 0.46 | 307.4 (26.7) | -27.3 | 0.01 | 0.67 |
| V̇O<sub>2peak</sub> (L·min<sup>-1</sup>) | 4.58 (0.38) | 8238.62 | 0.09 | 0.24 | 3.22 (0.28) | -5538.57 | 0.03 | 0.33 |
| V̇O<sub>2peak</sub> (mL·kg<sup>-1</sup>·min<sup>-1</sup>) | 53.5 (3.6) | -810.1 | 0.07 | 0.13 | 49.1 (4.3) | -922.6 | 0.17 | 0.03 |
| Rest. lactate concentration (mmol·L<sup>-1</sup>) | 1.42 (0.39) | 165.73 | 0.00 | 0.98 | 1.16 (0.29) | -667.26 | 0.00 | 0.94 |
| Max. lactate concentrations (mmol·L<sup>-1</sup>) | 10.6 (2.6) | 1077.4 | 0.06 | 0.16 | 10.0 (2.8) | 123.0 | 0.00 | 0.91 |
| Power output at VT1 (W) | 221.4 (44.6) | 6.7 | 0.00 | 0.88 | 149.2 (32.8) | -31.0 | 0.01 | 0.77 |
| V̇O<sub>2</sub> at VT1 (L·min<sup>-1</sup>) | 2.91 (0.47) | -718.34 | 0.00 | 0.87 | 1.91 (0.32) | -4040.93 | 0.02 | 0.70 |
| V̇O<sub>2</sub> at VT1 (mL·kg<sup>-1</sup>·min<sup>-1</sup>) | 33.9 (5.0) | -796.9 | 0.11 | 0.10 | 29.1 (4.9) | -548.8 | 0.08 | 0.20 |
| Power output at VT2 (W) | 343.6 (40.6) | 82.3 | 0.09 | 0.14 | 251.6 (25.7) | -84.4 | 0.05 | 0.16 |
| V̇O<sub>2</sub> at VT2 (L·min<sup>-1</sup>) | 4.11 (0.41) | 7254.77 | 0.07 | 0.17 | 2.90 (0.28) | -7904.09 | 0.06 | 0.08 |
| V̇O<sub>2</sub> at VT2 (mL·kg<sup>-1</sup>·min<sup>-1</sup>) | 47.9 (3.2) | -950.9 | 0.06 | 0.08 | 44.2 (4.4) | -966.8 | 0.19 | 0.01 |
| HR recovery 1 min (bpm) | -19.9 (6.0) | 313.2 | 0.03 | 0.23 | -22.3 (8.1) | 241.1 | 0.05 | 0.27 |
| HR recovery 3 min (bpm) | -18.7 (4.0) | 617.5 | 0.05 | 0.24 | -20.1 (3.1) | 678.1 | 0.05 | 0.14 |
| Peak power output (W) | 1166.7 (300.6) | 7.4 | 0.04 | 0.44 | 635.7 (162.0) | -5.1 | 0.01 | 0.67 |
| Mean power output (W) | 421.6 (42.7) | 15.2 | 0.00 | 0.71 | 282.4 (30.9) | -78.7 | 0.07 | 0.07 |
| Critical power (W) | 315.2 (48.4) | -48.2 | 0.04 | 0.25 | 224.3 (32.5) | -50.0 | 0.03 | 0.27 |
| W' (kJ) | 18.5 (6.5) | 482.7 | 0.08 | 0.06 | 10.5 (4.7) | -104.9 | 0.00 | 0.75 |
| V̇O<sub>2peak</sub> (L·min<sup>-1</sup>) | 4.51 (52.68) | 6261.2 | 0.07 | 0.22 | 3.16 (0.30) | -9437.27 | 0.10 | 0.04 |
| V̇O<sub>2peak</sub> (mL·kg<sup>-1</sup>·min<sup>-1</sup>) | 0.4 (4.8) | -539.5 | 0.06 | 0.18 | 48.2 (4.6) | -1054.57 | 0.26 | 0.00 |
| **Strength** |  |  |  |  |  |  |  |  |
| Leg iso. (N) | 4852 (1113) | -0.09 | 0.00 | 0.97 | 3621 (843) | -2.94 | 0.08 | 0.05 |
| Leg RFD 150 (N·s<sup>-1</sup>) | 14 (4) | -200.29 | 0.01 | 0.64 | 8 (3) | 934 | 0.08 | 0.29 |
| Leg con. [120 mm·s<sup>-1</sup>] (N) | 3504 (796) | 3.38 | 0.06 | 0.23 | 2609 (572) | -7.26 | 0.19 | 0.00 |
| Trunk iso. ext. (N) | 4223 (842) | 6.83 | 0.27 | 0.02 | 2283 (657) | -1.01 | 0.01 | 0.75 |
| Trunk iso. flex. (N) | 1995 (463) | -0.56 | 0.00 | 0.90 | 1153.90 (272.93) | -6.35 | 0.04 | 0.52 |
| Trunk con. ext. (N) | 4198 (607) | 3.04 | 0.03 | 0.27 | 2402 (432) | -0.88 | 0.00 | 0.80 |
| Trunk con. flex. (N) | 1826 (338) | -0.30 | 0.00 | 0.96 | 1227 (188) | -0.21 | 0.00 | 0.98 |
| CMJ height (m) | 0.41 (0.06) | -49413.77 | 0.07 | 0.13 | 0.31 (0.04) | -60457.51 | 0.08 | 0.18 |
| MTP iso. (N) | 3020 (555) | 9.10 | 0.22 | 0.04 | 2044.88 (444.06) | 1.31 | 0.00 | 0.81 |
| MTP RFD 150 (N·s<sup>-1</sup>) | 16 (6) | 634.26 | 0.09 | 0.17 | 12.24 (3.09) | -234.88 | 0.01 | 0.70 |
| MTP RFD 250 (N·s<sup>-1</sup>) | 21 (5) | 1035.59 | 0.22 | 0.01 | 14.52 (2.65) | 184.37 | 0.00 | 0.85 |
| Max GF (N) | 587 (101) | 62.69 | 0.34 | 0.03 | 370.39 (100.17) | -2.73 | 0.00 | 0.87 |
| Benchmarks |  |  |  |  |  |  |  |  |
| Back Squat (kg) | 178.3 (17.7) | -210.3 | 0.16 | 0.02 | 115.0 (13.8) | -267.2 | 0.17 | 0.00 |
| Deadlift (kg) | 211.6 (19.1) | -141.6 | 0.09 | 0.09 | 138.3 (16.1) | -252.2 | 0.18 | 0.18 |
| Bench Press (kg) | 127.9 (17.9) | -22.0 | 0.00 | 0.81 | 72.5 (9.2) | -218.2 | 0.05 | 0.26 |
| Snatch (kg) | 106.8 (10.6) | -317.0 | 0.13 | 0.03 | 71.3 (9.7) | -298.9 | 0.11 | 0.04 |
| Clean and Jerk (kg) | 132.2 (11.9) | -316.2 | 0.15 | 0.01 | 88.6 (10.1) | -443.7 | 0.24 | 0.00 |

Data are mean (standard deviation). Abbreviations: SD, standard deviation; CI, confidence interval; $\dot{V}\text{O}_{2\text{peak}}$ , peak oxygen consumption; VT1, first ventilatory threshold; VT2, second ventilatory threshold; max, maximal; W', work capacity above critical power; HR, heart rate; iso, isometric; RFD, rate of force development; con, concentric; ext, extension; flex, flexion; CMJ, counter movement jump; ext, extension; flex, flexion; con, concentric; CMJ, counter movement jump; MTP, mid-thigh pull; GF, grip force.

Moreover, an assessment of explosivity, as measured by the height in the countermovement jump, was strongly positively associated with performance in both sexes. Interestingly, absolute strength, as assessed with the mid-thigh pull, had a slight negative association with Open performance in men, suggesting that stronger performance in this parameter could potentially impair Open rankings.

Lastly, we found that sport-specific tests, including compound lifts, as well as weightlifting exercises such as clean and jerk and snatch, showed strong evidence for an association with Open performance. However, it is important to note that these results are based on self-reported data, unlike the other parameters discussed earlier, which were measured under highly controlled laboratory conditions.

The coefficients presented in Table 3 are practically applicable for estimating potential improvements in CrossFit^®^ Open performance when a single parameter is enhanced or altered. For instance, a shorter core length by 10cm is associated with a loss of 17’719 rank points for men, translating to an approximate improvement in rank by 4’860 positions. Similarly, a 10 cm shorter arm length was associated with 15’609 less rank points among men, improving an athlete by approximately 4’320 ranks and vice versa. When considering trainable parameters, an improved 1RM Clean and Jerk by 20kg for example is associated with a loss of 8’874 rank points among female athletes, placing them approximately 2’280 ranks higher on the leaderboard worldwide.

## DISCUSSION

The primary objective of this study was to establish a comprehensive profile for laboratory-assessed endurance and strength parameters in highly-trained CrossFit^®^ athletes. This was done with an aim to identify parameters that could reliably predict overall CrossFit^®^ performance. Our findings suggest that basic anthropometric data, including parameters like height, thigh length, or body weight, are highly predictive of performance in the CrossFit^®^ Open for both sexes. In contrast, standard laboratory endurance and strength tests appeared less predictive and showed more variability between men and women.

CrossFit^®^ competitions seek to identify the ‘fittest’ individuals based on their ability to show increased work capacity across broad time and modal domains.^18^ With the emergence of CrossFit^®^ as a competitive sport, multiple exercise modalities are combined at high intensity with performance as the critical metric. This approach epitomizes the concept of concurrent training, blending elements of strength and endurance training, as well as skill and agility.

Over the years, numerous studies have strived to characterize CrossFit^®^ athletes, focusing on pinpointing the key performance indicators crucial for success in CrossFit^®^-specific workouts. The primary concentration of these studies has been on sport-specific tasks typically included in standard CrossFit^®^ training programs such as barbell back squats and ergometer time trials. Despite the useful insights, their emphasis on trainable variables, which inherently and automatically correspond with Open performance, might provide an incomplete representation of the true determinants of success in CrossFit^®^. Thus, this study utilized a battery of gold-standard laboratory-based tests, a novel approach providing an unbiased understanding of what is essential for superior CrossFit^®^ performance. Our study participants included athletes in the top 5% of the CrossFit^®^ Opens between 2019-2022, a cohort that represents a highly trained segment of CrossFit^®^ competitors, including Semi-finalists and Games competitors. Intriguingly, linear regression analysis revealed a multitude of anthropometrical parameters with strong correlations to CrossFit^®^ Open performance. This suggests a noticeable advantage for smaller athletes with shorter limbs in the Open competition. As CrossFit^®^ is strongly based on weightlifting movements it can be expected that athletes with shorter limbs have an advantages as it was previously described for Olympic weightlifting ^19^ and power lifting.^20^ The potential rationale behind this advantage are the decreased mechanical torque required to move a given weight by smaller athletes and athletes with smaller limbs.^19^ Further the amount of muscular work required is decreased via the smaller vertical distance that the barbells needs to be moved.^19^ Both these aspects likely lead to more efficient (energy-saving) and faster completion of Olympic weightlifting and other barbell, dumbbell, kettlebell or even odd object movements by shorter athletes. This advantage is likely pronounced when these exercises are performed over multiple reps as often seen in CrossFit^®^ Open workouts. The potential disadvantages for taller athletes regarding longer levers, higher required torque and larger distances to cover likely also hold true for bodyweight (gymnastics) movements such as pull-ups or push-ups.^19^ Taller athletes, due to their anthropometry, must therefore generate higher power outputs to complete the same workouts, potentially placing them at a disadvantage.

This suggests that the CrossFit^®^ Open may not provide an equally competitive platform for all athletes. While it is potentially true that final stages of competition (Semi-finals and Games), may offer more balanced challenges, this aspect was not investigated in the present study. To address this potential bias, it might be beneficial to incorporate more exercises and movements into the CrossFit^®^ Open workouts where taller athletes naturally have an advantage, such as rowing, box jumps, rope climbs or wall balls.

The primary objective for competitive CrossFit^®^ athletes is to demonstrate exceptional capabilities across diverse exercise modalities and throughout various time domains. This objective is distinctly different from specialized sports such as cycling, long-distance running, or weightlifting. To better comprehend where an elite CrossFit^®^ athlete stands in terms of strength, explosivity, endurance capacity, and threshold power relative to a specialist in their respective field, we utilized data from past studies on elite male and female cyclists and weightlifters.^6–10^ This allowed us to compare our cohort’s lab-based parameters with those of specialists. Furthermore, we differentiated between our well-trained cohort and our elite cohort (those reaching semi-finals and beyond).

On the whole, elite CrossFit^®^ athletes performed on average 10.8% less than their specialist counterparts. V□O_2peak_ was notably lower in our group, averaging a modest 54.7 mL·kg^-1^·min^-1^ in male athletes, versus reported 76.8 mL·kg^-1^·min^-1^ in male elite cyclists.^6^ In female CrossFit^®^ athletes the same observation was made with 50.15 mL·kg^-1^ min^-1^ versus reported 66.6 mL·kg^--1^·min^-1 6^. It is quite intriguing that this group of highly trained CrossFit athletes displayed a rather low relative V□O_2peak_. The observed relative V□O_2peak_ values of CrossFit^®^ athletes in our study were 15% (53.7 vs 46.6 mL·kg^-1^·min^-1^) and 24% (48.6 vs 39.3 mL·kg^-1^·min^-1^) above values reported of not specifically trained healthy individuals with a comparable average age and from the same geographic region.^21^ CrossFit^®^ athletes typically possess high relative muscle mass and BMI, which could contribute to the comparatively low relative V□O_2peak_ identified in this study. Other parameters, such as mid-thigh pull strength and to a lesser extent, countermovement jump height, were also considerably lower than those of specialists, as indicated by Figure 3a and 3b. This further underscores that CrossFit^®^ athletes do not specialize in a single discipline and that concurrent training can compromise performance at the extremities of the physiological spectrum towards both endurance and strength or explosivity. This study specifically recruited and included the same number of male and female athletes to address the lack of data for women in performance sport and to display any potential notable sex differences. Performance metrics differed between men and women from 11% for relative V□O_2peak_, 29% for countermovement jump, 42% for isometric leg strength, 42% for critical power, 45% for absolute V□O_2peak_, 55% for mid-thigh-pull and 100% for maximum work above critical power (W’) which are all similar to sex differences observed in specialists.^6–10,22^ As expected, performance metrics not normalized to bodyweight showed larger sex differences then metrics normalized to bodyweight. Correlations between cardiorespiratory fitness metrics and CrossFit^®^ Open performance in women, along with a relatively small sex difference in relative V□O_2peak_, suggest greater importance of cardiorespiratory fitness for females in CrossFit^®^. Anthropometric characteristics, however, hold more significance for males, possibly due to larger variations in these markers among male subjects. Optimal anthropometric characteristics appear comparable for both sexes, given the similar test conditions, except for around 30% lower weights in Open workouts for women. More male study participants exhibit suboptimal body height, limb length (both to tall) and weight (too heavy) whereas females tend to align naturally with more optimal anthropometric characteristics, thus playing a minor role for women.

**Figure 3a.**
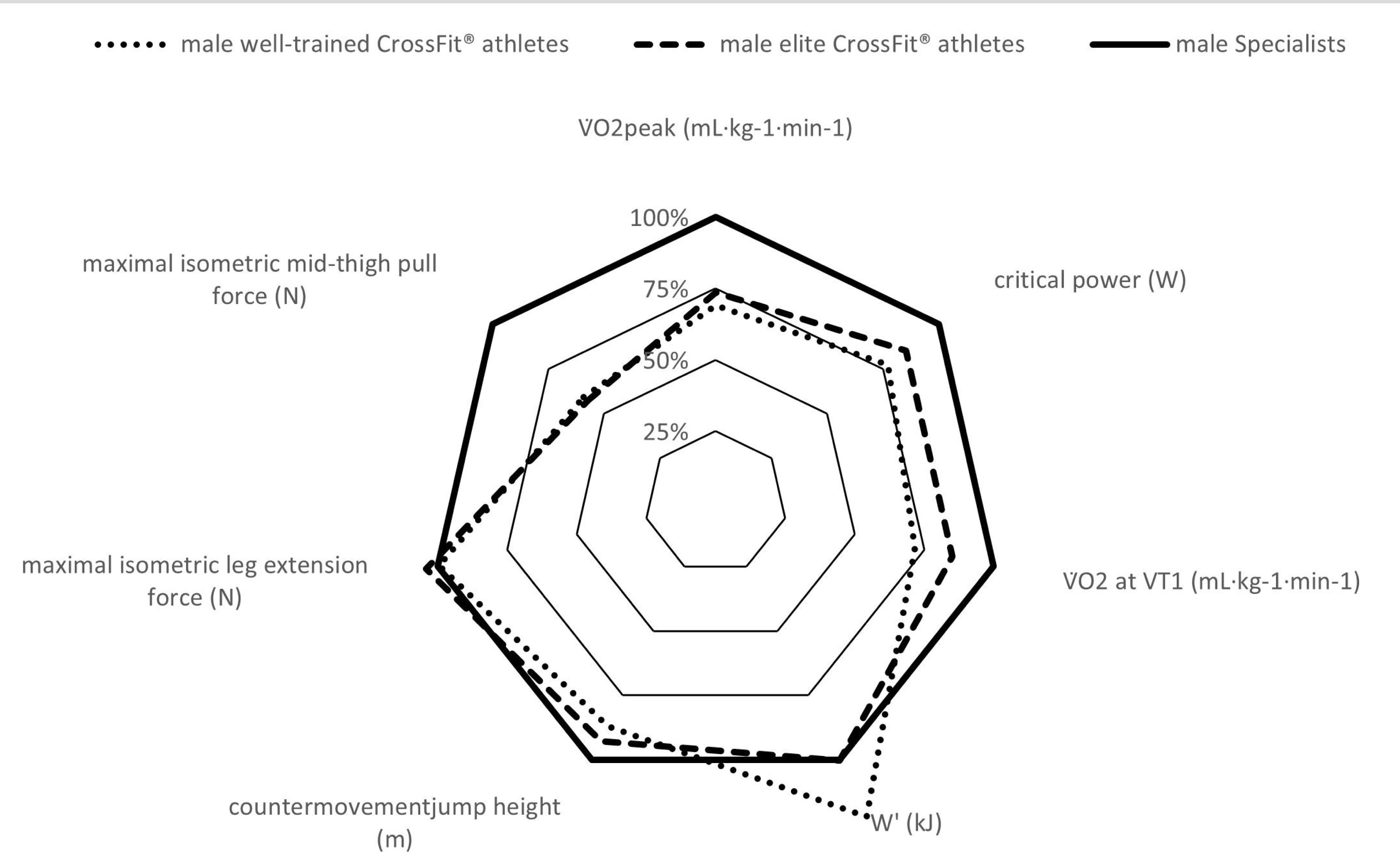
Comparison of male well-trained CrossFit^®^ athletes (all top 5% CrossFit^®^ athletes without athletes reaching the semi-finals) versus male elite CrossFit^®^ athletes (reaching at least the semi-finals) versus male elite specialized athletes on world championship/ world cup level (for cardiopulmonary parameters cyclists^6,7^ and for strength parameters weightlifters^8–10^). Data are CrossFit^®^ athlete’s mean relative to specialized athlete’s mean in percentages. Specialized athletes are 100%. Abbreviations: VDO_2peak_, peak oxygen consumption; VT1, first ventilatory threshold; W’, work capacity above critical power.

**Figure 3b.**
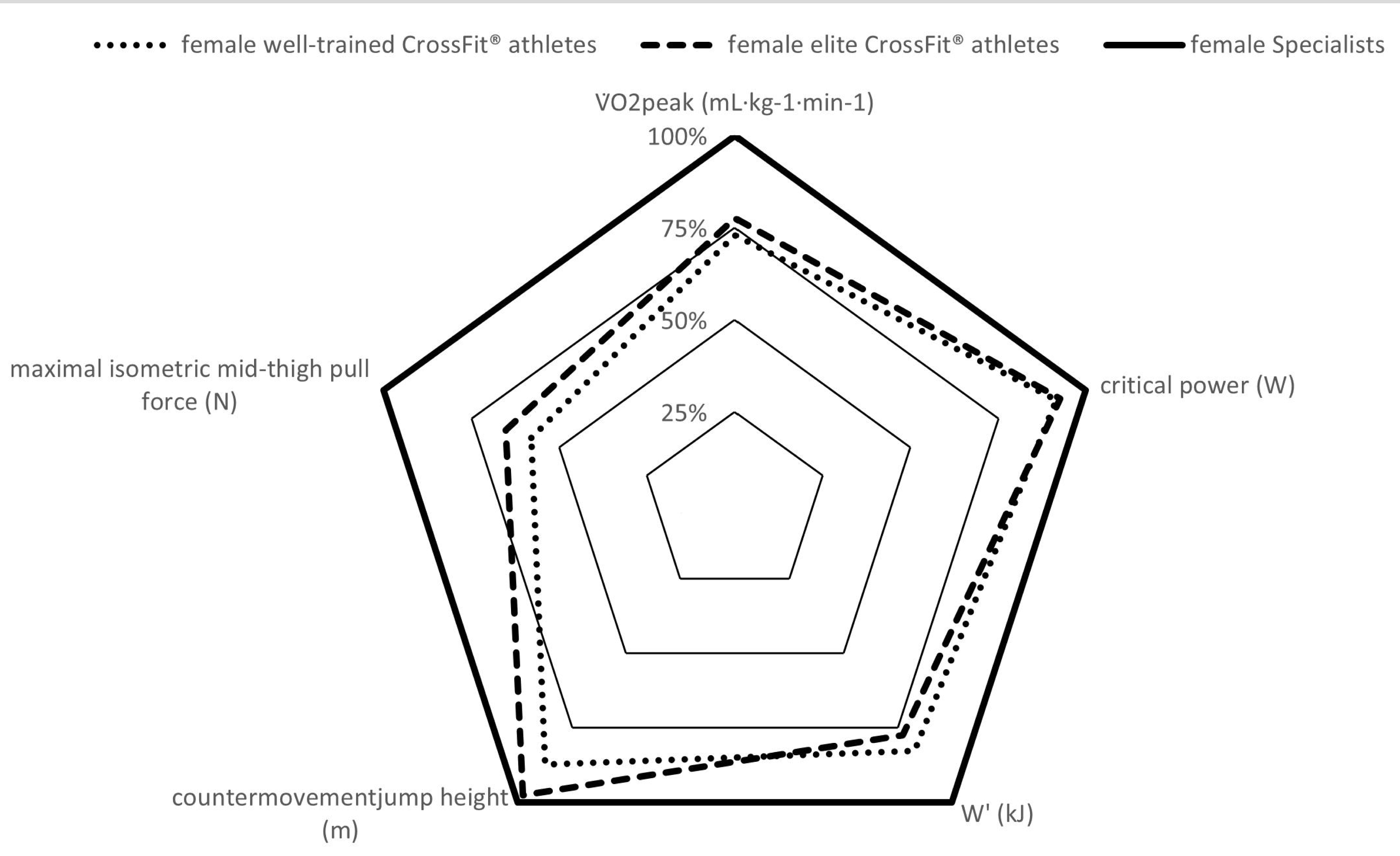
Comparison of female well-trained CrossFit^®^ athletes (all top 5% CrossFit athletes without athletes reaching the semi-finals) versus female elite CrossFit^®^ athletes (reaching at least the semi-finals) versus female specialized athletes on world championship/ world cup level (for cardiopulmonary parameters cyclists^6^ and for strength parameters weightlifters^8,22^). Data are CrossFit^®^ athlete’s mean relative to specialized athlete’s mean in percentages. Specialized athletes are 100%. Abbreviations: VDO_2peak_, peak oxygen consumption; W’, work capacity above critical power.

### Practical applications

This study critically enhances both the theoretical knowledge and practical comprehension of CrossFit^®^, a unique and relatively novel sport. Uncovering the fundamental physiological parameters and CrossFit^®^-specific benchmarks, it delivers crucial insights for athletes and coaches to optimize their training regimens. It highlights the physiological benchmarks in terms of endurance capacity and strength that need to be achieved to make it to the highest level in this sport. Notably, the study underscores the essential role of anthropometrics in CrossFit^®^, evidencing a competitive edge for shorter athletes with shorter limbs in the Open competitions. As such, if the ultimate aim of CrossFit^®^ is to crown the fittest individual, regardless of anthropometric characteristics, our data recommend considering future adjustments to CrossFit^®^ Open workout designs.

Future research should aim to track athletes across multiple competitive seasons, analyzing physiological markers linked to improvements or deteriorations not only in the CrossFit^®^ Open, but also in the Quarterfinals, Semi-finals, and the CrossFit^®^ Games. Incorporating on-site testing during Semi-finals or the Games could simplify recruiting and enhance our understanding of the physiological characteristics of top athletes in this sport.

## CONCLUSION

In conclusion, this study established a comprehensive profile of endurance, strength and anthropometric parameters in high-performing CrossFit^®^ athletes. The study revealed evidence for an association between anthropometric parameters, specifically leg and thigh length and CrossFit^®^ Open performance for male and female and emphasize the advantage of athletes with shorter legs and thighs. Standard laboratory performance tests displayed less predictive power and greater variability between men and women. Elite CrossFit^®^ athletes demonstrated approximately 10% lower performances than specialists in specific disciplines but showed high numbers across a broad range of endurance and strength parameters.

The study informs training optimization and suggests potential modifications to Open workout designs to level the playing field for athletes across different anthropometrics characteristics.

## Acknowledgments

We thank all the participants who made this study possible by their participation.

Conflict of interest:

None of the authors involved in the present study have any conflict of interest, financial, personal, or otherwise, which would influence this research. The authors declare that the results of the study are presented clearly, honestly, and without fabrication, falsification, or inappropriate data manipulation.

## Authors’ contributions

D.H., G.H. and J.W. wrote the original draft of the article. R.L, J.A, E.W., R.R., A.S.T., and R.K. participated in the review and editing of the article. D.H. did the visualization of the data. G.H., R.K., and JW. participated in the concept and design. R.L., J.A., E.W. and J.W. participated in the data acquisition. D.H., G.H., and J.W. participated in the data analysis and interpretation.

## References

1. CrossFit - The Official Website. Accessed June 23, 2023. https://games.crossfit.com/history-of-the-games

2. Mangine GT, Tankersley JE, Mcdougle JM, et al. Predictors of CrossFit Open Performance. Sports. 2020;8(7):102. doi:10.3390/sports8070102

3. Martínez-Gómez R, Valenzuela PL, Alejo LB, et al. Physiological Predictors of Competition Performance in CrossFit Athletes. International Journal of Environment Research and Public Health. 2020;17(10):3699. doi:10.3390/ijerph17103699

4. Butcher S, Neyedly T, Horvey K, Benko C. Do physiological measures predict selected CrossFit® benchmark performance? Open Access J Sports Med. Published online July 2015:241. doi:10.2147/oajsm.s88265

5. Bellar D, Hatchett A, Judge LW, Breaux ME, Marcus L. The relationship of aerobic capacity, anaerobic peak power and experience to performance in CrossFit exercise. Biol Sport. 2015;32(4):315–320. doi:10.5604/20831862.1174771

6. Almquist NW, Hansen J, Rønnestad BR. Development of Cycling Performance-Variables and Durability in Female and Male National Team Cyclists: From Junior to Senior. Med Sci Sports Exerc. Published online 2023. https://journals.lww.com/acsm-msse/Fulltext/9900/Development_of_Cycling_Performance_Variables_and.297.aspx

7. Lucía J; Durántez A; Hoyos J; Chicharro J L AP. Physiological Differences Between Professional and Elite Road Cyclists. Int J Sports Med. 1998;19(05):342-348. doi:10.1055/s-2007-971928

8. Stone MH, Sands WA, Pierce KC, Carlock J, Cardinale M, Newton RU. Relationship of maximum strength to weightlifting performance. Med Sci Sports Exerc. 2005;37(6):1037–1043. doi:10.1249/01.mss.0000171621.45134.10

9. Hiikkinen K, Komi P V, Alan M, Kauhanen H. EMG, muscle fibre and force production characteristics during a 1 year training period in elite weight-lifters. Eur J Appl Physiol Occup Physiol. 1987;56:419–427.

10. Zaras N, Stasinaki AN, Spiliopoulou P, Arnaoutis G, Hadjicharalambous M, Terzis G. Rate of Force Development, Muscle Architecture, and Performance in Elite Weightlifters. Int J Sports Physiol Perform. 2021;16(2):216–223. doi:10.1123/ijspp.2019-0974

11. Shephard RJ. PAR-Q, Canadian Home Fitness Test and Exercise Screening Alternatives. Sports Medicine. 1988;5(3):185–195. doi:10.2165/00007256-198805030-00005

12. Maier T. Manual Leistungsdiagnostik. Swiss Olympic; 2016.

13. Matheson LA, Duffy S, Maroof A, Gibbons R, Duffy C, Roth J. Intra- and inter-rater reliability of jumping mechanography muscle function assessments. Journal of musculoskeletal & neuronal interactions. 2013;13(4):480—486. http://europepmc.org/abstract/MED/24292618

14. Ruiz-Ruiz J, Mesa JLM, Gutiérrez A, Castillo MJ. Hand size influences optimal grip span in women but not in men. Journal of Hand Surgery. 2002;27(5):897–901. doi:10.1053/jhsu.2002.34315

15. Borg G. Borg’s Perceived Exertion And Pain Scales.; 1998.

16. Vanhatalo A, Doust JH, Burnley M. Determination of critical power using a 3-min all-out cycling test. Med Sci Sports Exerc. 2007;39(3):548–555. doi:10.1249/mss.0b013e31802dd3e6

17. Jones AM, Vanhatalo A, Burnley M, Morton RH, Poole DC. Critical power: Implications for determination of ⊙ O2max and exercise tolerance. Med Sci Sports Exerc. 2010;42(10):1876–1890. doi:10.1249/MSS.0b013e3181d9cf7f

18. Glassman G. WHAT IS FITNESS? The CrossFit Journal. 2002;(2). Accessed June 26, 2023. http://library.crossfit.com/free/pdf/CFJ-trial.pdf

19. Storey A, Smith HK. Unique Aspects of Competitive Weightlifting. Sports Medicine. 2012;42(9):769–790. doi:10.1007/BF03262294

20. Cooke DM, Haischer MH, Carzoli JP, et al. Body Mass and Femur Length Are Inversely Related to Repetitions Performed in the Back Squat in Well-Trained Lifters.; 2019. www.nsca.com

21. Wagner J, Knaier R, Infanger D, et al. Novel CPET Reference Values in Healthy Adults: Associations with Physical Activity. Med Sci Sports Exerc. 2021;53(1):26–37. doi:10.1249/MSS.0000000000002454

22. Haugen TA, Breitschädel F, Wiig H, Seiler S. Countermovement Jump Height in National-Team Athletes of Various Sports: A Framework for Practitioners and Scientists. Int J Sports Physiol Perform. 2021;16(2):184–189. doi:10.1123/ijspp.2019-0964

