## Supplemental material for "Physiological Profiles of Male and Female CrossFit^®^ Athletes"

*Supplemental material to an article in International Journal of Sports Physiology and Performance*

**MATERIAL AND METHODS**

**Acquisition of participant characteristics.**

Extremities' lengths were measured in a standardized way (see Table S1).

Table S1. Extremities' measurement.

| **Extremity** | **Anatomical fix points** |
| --- | --- |
| **Legs** | **FROM** Spina iliaca anterior superior **TO** Caput tibiae **TO** Most distal part of the lateral malleolus |
| **Thighs** | **FROM** Spina iliaca anterior superior **TO** Caput tibiae |
| **Lower Legs** | **FROM** Caput tibiae **TO** Distal part of the lateral malleolus |
| **Upper body** | **FROM** Lateral end of clavicula **TO** Spina iliaca anterior superior |
| **Arms** | **FROM** Acromion **TO** distal end of the radius |

Forced vital capacity (FVC) and forced expiratory volume in one second (FEV1), objective parameters of respiratory function, were measured in accordance with the American Thoracic Society/European Respiratory Society guidelines^1^ immediately before the cardiopulmonary exercise test.^2^

**Comprehensive overview of the exercise protocol**

**Strength testing.** Before strength testing, all participants performed a standardized warm-up program shown in Table S2.

Table S2: Standardized warm-up program

5 minutes warm-up on the rowing machine at moderate intensity (Borg 11 of 20).

20 squats, 4 sets of 12-10-8-6 squats with a 20 kg barbell, 30% 1RM, 40% 1RM and 50% 1RM

20 "Goodmornings" with the band or barbell, 15 "Deadlifts" and 15 "Cleans" with a 20 kg barbell

Before each measurement, practice trials were completed. For isometric strength assessment one practice trial was performed, for isokinetic movements one practice trial with 3 to 5 repetitions and for maximal force production of jump height, mid-thigh pull, and grip force two practice trials were undertaken. Isometric measurement consisted of three trials with one-minute rest (for legs, jump height, mid-thigh pull and grip force) or 30-second rest (for trunk) in between each trial. In all tests, the average of the two best trials was then included in the analysis. Isometric contraction had to be as "fast" and as "strong" as possible with the feet against the surface and to maintain the generated force for five seconds. Isokinetic measurements consisted of two trials, each with five repetitions and a one-minute rest (for legs) and two-minute rest (for trunk) in between. Only the better trial was included in the analysis. Participants were encouraged to press as “fast and strong as possible” to calculate rate of force development.^3^

**Mid-thigh pull strength testing.** The shoulders were above the bar, causing the upper body to be bent forward at 5-10°. The shoulder blades have been actively contracted in this position. The type of grip could be chosen individually. Tape for the hands and magnesia were allowed, but no traction aids.

Cardiopulmonary exercise testing. CPET was performed using a ramp protocol on a cycle ergometer. Before starting the test, the equipment was calibrated in standard fashion with reference gas and known volume. The cycle ergometer's seat height and handlebar setting were individually adjusted to the participants' physique. It was possible to use the participant’s own cycling shoes with a click system if available.

Participants were allowed to choose their pedaling cadence as long as it was maintained above 60 rpm. After a 5-min warm-up at 100 W for males and 50 W for females, the workload increased linearly with 30 W per min for all participants until maximal voluntary exhaustion was reached.

In addition to gas exchange measurement, heart rate was also measured continuously with a 12-lead electrocardiograph (Custo med GmbH, Ottobrunn, Germany). Participants' rating of perceived exertion was assessed according to the Borg scale 6-20.^4^ For gas exchange analysis, the data were averaged over 10-second intervals. Blood samples were taken at the earlobe before the start of the test, immediately after test termination, and after two, three, and five minutes into the recovery phase. Blood lactate values were measured using the Biosen C-line lactate analyzer (EKF-diagnostic GmbH, Barleben, Germany).

**3-minutes all-out test.** After the ramp test, a 20-min recovery (5 min active recovery at 30 W + 15 min passive recovery) was conducted. Participants were instructed to increase their cadence to approximately 110-120 rpm during the last 10 seconds of the 3-minute unloaded cycling. The unloaded phase was followed by 3MT, in which participants were required to rapidly increase their cadence with the intention of "cycling as fast as possible". To provide maximum exhaustion, participants were instructed to sustain their cadence throughout the 3-min duration of the test as high as possible. The resistance changes with changes in cadence according to the formula Workload (W)= α∙cadence^2^. A fixed resistance was calculated for each participant. The described procedure of the 3MT is well established and reported in detail in Constantini et al.^5^.

**Statistical Analysis.**

The 2022 CrossFit® Open consisted of three workouts (see Table S3).

**Table S3.** 2022 CrossFit Open Workouts

| **Workout** | **Modality** | **Content** |
| --- | --- | --- |
| **22.1** | As many repetitions as possible in 15min | 3 wall walks  12 dumbbell snatches  15 box jump-overs  (Men: 50-lb. dumbbell, 24-in. box;  Women: 35-lb. dumbbell, 20-in. box) |
| **22.2** | 1-2-3-4-5-6-7-8-9-10-9-8-7-6-5-4-3-2-1  Repetitions on time with a time limit of 10min | Deadlifts  Bar-facing Burpees  (Men: 255-lb. barbell;  Women: 155-lb. barbell) |
| **22.3** | On time with a time limit of 12min | 21 pull-ups; 42 double-unders; 21 thrusters (weight 1)  18 chest to bar pull-ups; 36 double-unders; 18 thrusters (weight 2)  15 bar muscle ups; 30 double-unders;15 thrusters (weight 3)  (Men: 95 lb (weight 1), 115 lb (weight 2), 135 lb (weight 3);  Women: 65 lb (weight 1), 75 lb (weight 2), 85 lb (weight 3)) |

**REFERENCES**

1.	Miller MR, Hankinson J, Brusasco V, et al. Standardisation of spirometry. *European Respiratory Journal*. 2005;26(2):319-338. doi:10.1183/09031936.05.00034805

2.	Balady GJ, Arena R, Sietsema K, et al. AHA Scientific Statement Clinician’s Guide to Cardiopulmonary Exercise Testing in Adults A Scientific Statement From the American Heart Association. Published online 2010. doi:10.1161/CIR.0b013e3181e52e69

3.	Maffiuletti NA, Aagaard · Per, Anthony ·, et al. Rate of force development: physiological and methodological considerations. *Eur J Appl Physiol*. 2016;116:1091-1116. doi:10.1007/s00421-016-3346-6

4.	Borg G. *Borg’s Perceived Exertion And Pain Scales*.; 1998.

5.	Constantini K, Sabapathy S, Cross TJ. A single-session testing protocol to determine critical power and W′. *Eur J Appl Physiol*. 2014;114:1153-1161. doi:10.1007/s00421-014-2827-8
